# Induction of tumor-initiating cells and glioma-initiating cells from fetal neural stem cells through p53 genome editing

**DOI:** 10.1101/2022.02.24.481795

**Authors:** Naoki Ohtsu

## Abstract

The elucidation of the properties of malignant glioma and development of therapeutic methods require model cells with characteristics such as invasiveness, multinuclearity, and ability for mitosis. A previous study has shown that overexpression of active HRas (HRasL61) in neural stem/progenitor cells isolated from fetal telencephalic neuroepithelial cells with homozygous deletion of the p53 locus forms glioma-initiating cells (GICs). The orthotopically transplantation of 10 cells into the forebrain of immunodeficient mice, resulted in the development of glioblastoma multiforme-like malignant brain tumors. In this study, GICs were induced from wild-type fetal neural stem/progenitor cells. Through forced expression of CRISPR-associated protein 9 together with a guide RNA that recognizes the p53 gene, neural stem/progenitor cells containing p53 knockout cells were obtained. Furthermore, HRasL61 was forced-expressed and transplanted into the immunodeficient mice. These cells were tumor-initiating cells (TICs) with tumorigenicity in the brain. However, no invasiveness to the brain was observed. The p53 homozygous-deleted cells obtained through single-cell cloning from TICs were GICs, forming glioblastoma multiforme-like tumors. The induced GICs strongly expressed the GIC marker. These results indicate that GICs can be induced through genetic recombination using genome editing. This study suggests that the use of genome editing may lead to the elucidation of the oncogenic mechanisms of human cells that cannot delete genes.

## Introduction

Glioblastoma multiforme (GBM; WHO grade IV) is a malignant brain tumor with an average survival time of proximately 15 months, and a high probability of recurrence.

The GBM tumor is a heterogeneous cell population, which contains highly invasive, drug-resistant and radioresistant cell populations. These cells are termed glioma-initiating cell (GIC) [1]. To analyze the characteristics of GICs in GBM, human GIC strains have been established using tumors from GBM patients and GBM-like artificial GIC (induced GIC: iGIC) strains have been established from mouse NSCs [2–5]. The concept of inducible cancer-initiating cells was proposed, triggered by the transformation into cells with high tumorigenicity through the introduction of the three factors Ras, IkB and c-Myc into differentiated epithelial cells [6]. In the preparation of iGICs, fetal neural stem cells (NSCs) of p53 knockout (KO) mice were overexpressed with activated HRas (HRasL61) [2]. The Ras signaling pathway is involved in human gliomas, (i.e., 90% of multiple glioblastomas) [7–9]. p53 is the most frequently mutated tumor suppressor gene in GBM [7,8,10–12]. Forced expression of HRasL61 in p53 KO NSCs induced GICs to form GBM-like tumors exhibiting invasiveness, multicellularity, pleomorphism, multinucleated giant cells, mitosis, and necrosis. It has also been reported that GICs are induced by the forced expression of HRasV12 in the NSCs of Arf/INK4a locus null mice [13]. An investigation of genes specifically expressed in GBM, analysis of gene function, and analysis of the mechanism of action have been performed using GIC [2–4,14,15].

In cancer research, genome editing technology using the Crispr/Crispr-associated protein 9 (Cas9) system has been widely used in recent years, leading to the development of new gene therapies [16]. The production of GICs through genome editing may eliminate the need to bread KO mice.

Patient-derived xenograft experiments, in which human GIC strains established from GBM patients were transplanted into the brains of immunodeficient mice, have been performed. Some strains were tumorigenic, however, they produced tumors outside the brain rather than in the brain. It was necessary to clarify the possibility of technical problems and the potential properties of these cells.

In this study, we describe a new establishment method for iGICs using genome editing and compare the gene expression with induced tumor-Initiating Cell (TIC) strains, also established during this process.

## Materials and Methods

### Animals

C57BL/6 mice (1 pregnant mouse and 12 female mice) and NOD-SCID mice (25 female mice) were purchased from CLEA Japan (Tokyo, Japan). Mice were maintained under specific-pathogen free conditions with temperature, humidity and lighting controlled by authors and experienced expertise at Laboratory of Animal Experiment, Institute for Genetic Medicine, Hokkaido University. The mice were housed in plastic cages with wire mesh lids and water bottles (CLEA Japan, Tokyo, Japan), and were put on the open racks. Mice were fed with Labo MR Stock diet (Nosan, Kanagawa, Japan) and maintained on wood chip bedding (Iwakura, Hokkaido, Japan). All animal experiments were conducted in accordance with the Hokkaido University Guide for the Regulation on Animal Experimentation. All experiments were reviewed and approved by the Animal Care and Use Committee of the Hokkaido University (Permit Number: 16-0003). All animals’ health condition and behavior were monitored every day. Prior to our experimental endpoint, no mice became severely ill or died, or display any abnormality. Humane endpoints were determined by manifestation of neurological signs or 30 days post transplantation within our experiments.

When reaches humane endpoint, all mice were euthanized through cervical dislocation performed by trained individuals using appropriate equipment. Within our 1-year experiments, all mice were euthanized when reaches humane endpoint, or 30 days post-transplantation. All surgery was performed under 10% pentobarbital anesthesia, and all efforts were made to minimize suffering and distress, and stress.

### Cell culture

Pregnant mouse was euthanized by cervical dislocation and embryonic day14.5 mouse embryos were obtained. Mouse NSCs were prepared from the telencephalon of mice embryo [17]. NSCs were expanded and cultured in Dullbecco’s modified Eagle’s medium/Nutrient Mixture F-12 (SIGMA) containing bovine insulin (10 μg/mL), human apo-transferrin (100 μg/mL), bovine serum albumin (100 μg/mL), progesterone (60 ng/mL), putrescine (16 μg/mL), sodium selenite (40 ng/mL), N-acetylcysteine (60 μg/mL), forskolin (5 μM), basic fibroblast growth factor (10 ng/mL), epidermal growth factor (10 ng/mL), penicillin, and streptomycin (Nakarai). NSCL61 was cultured in a half-medium containing a 1:1 mixture of NSC medium and Dullbecco’s modified Eagle’s medium with 10% fetal calf serum (HyClone), penicillin, and streptomycin (fetal calf serum medium), as described previously described [5]. Cells were cultured at 37°C and under 5% carbon dioxide.

### Construction of the genome editing vector

The following single-guide RNA targeting p53 and luciferase as control (CTRL) were designed using the CRISPR direct online tool (http://crispr.dbcls.jp/). The sequences of oligonucleotide for single-guide RNA are showed in S1 Table. It was annealed to the complementary oligonucleotide and cloned into pX330A-1×2 (Plasmid #58766, addgene) [18]. DNA sequencing was performed using an ABI 3130xl genetic analyzer (Life Technologies) with a Big Dye Terminator v3.1 cycle sequencing kit (Life Technologies).

### Transfection

The pX330A-1×2 containing oligo DNA was transfected into NSCs and HRasL61 [2] containing pCMS-HRasL61-EGFP was transfections into gp53-NSCs using a CUY21 SC Square Wave Electroporator according to the manufacturer’s instructions (Neppagene). Briefly, 1×10^6^ cells were suspended in 100 μL Opti-MEM (Thermo Fisher Scientific) with 10 μg vectors and subsequently transfected using the electroporator. Green fluorescent protein (GFP) positive cells in HRasL61 with GFP transfected cells were sorted using flow-cytometry (FACS Aria II; BD Bioscience).

### Genomic polymerase chain reaction

Genomic DNA was extracted from the gp53-NSCL61 clone, generated tumor, and recipient mice using proteinase K (Roche)-containing buffer. Subsequently, phenol/chloroform isolation and ethanol precipitation were performed. Genomic polymerase chain reaction (PCR) was performed using Taq Polymerase (Jena Bioscience) with the primers listed in S2 Table. The reaction conditions were 95°C for 1 min, 40 cycles at 95°C for 20 s, 60°C for 30 s, and 72°C for 1 min. The PCR amplicons were analyzed with electrophoresis in a 1% agarose gel.

### Western blotting

Western blotting was performed as previously described [3]. The blotted membranes were probed with anti-p53 antibody (1:500; Leica Biosystems) and anti-GAPDH antibodies (1:1000, ThermoFisher Scientific), and the signals were detected by enhanced chemiluminescence and Chemidock MP (Bio-Rad).

### T7 Endonuclease1 assay

The target site of genomic DNA was amplified with PCR using the appropriate primer set (see S1 Table). The condition of the first reaction were 95°C for 1 min, 40 cycles at 95°C for 20 s, 60°C for 30 s, and 72°C for 1 min. The PCR amplicon was purified using a FastGene gel extraction kit (Nippon Genesis). The hybridization conditions were 95°C for 5 min, 95°C to 85°C (−2°C/s), 85°C to 25°C (−0.1°C/s), and finally 4°C for cooling. This fragment was digested with T7E1 Endonuclase1 (New England Biolabs) for 15 min at 37°C. The products were analyzed through electrophoresis in a 1% agarose gel.

### Intracranial cell transplantation into NOD-SCID mice brains and brain-tumor histopathology

CTRL-NSCL61 and gp53-NSCL61s were suspended in culture medium and injected into the brains of 5-8-week-old female mice, as previously described [14,19]. Mice were maintained until manifestation of neurological signs or for 30 days post-transplantation. Mice were then anesthetized and underwent cardiac perfusion with 4% paraformaldehyde. The mouse brains were dissected, fixed in 4% paraformaldehyde overnight, transferred to 70% ethanol, processed on Tissue-Tek VIP (Sakura Finetek Japan), and embedded in paraffin. Coronal sections (6-μm thick) from the cerebral cortex were prepared on a microtome and stained with hematoxylin eosin using standard techniques. Sections were observed using all-in-one fluorescence microscopy (BZ-X800; KEYENCE) and its image joint software.

### Immunohistochemistry

Paraffin-sections were hydrolyzed after deparaffinization. After washing with phosphate-buffered saline (PBS), the sections were permeabilized with 0.3% Triton X-100 in PBS for 10 min at room temperature. The sections were subsequently pretreated with 5% skim milk in PBS-0.3% Triton X-100 for 30 min at room temperature, and incubated with rabbit anti-GFP antibody (1:200; MLB) for 16 h at 4°C. The antibody used was the Alexa488-conjugated goat anti-rabbit IgG (1:1,000; Invitrogen). The sections were counterstained with 4’,6-diamidino-2-phenylindole (DAPI, 1 μg/mL) to label the nuclei and were observed using all-in-one fluorescence microscopy (BZ-X800; KEYENCE).

### Quantitative reverse transcription PCR

Total RNA was purified from wild-type NSCs (WT-NSCs) and gp53-NSCL61s using the Trizol reagent (Invitrogen). In all, 1 μg of total RNA was used for quantitative reverse transcription reaction-PCR (RT-qPCR) with ReverTra Ace (Toyobo), according to the manufacture’s instructions. qPCR was performed with the THUNDERBIRD SYBR qPCR Mix (Toyobo) and analyzed using the StepOne real-time PCR system (Applied Biosystems). The primer sequences are shown in S3 Table.

### Statistical analyses

Data are shown as mean ±standerd deviation. The Student’s t-test was used for analysis. All statistical analyses were performed with Excel 2019 (Microsoft). The survival data were analyzed for significance through the Kaplan-Meier method using GraphPad Prism version 4 software. The p-values were calculated using the log-rank test.

## Results

### Generation of gp53-NSCL61 from WT fetal mice NSCs using genome editing

The CRISPR/Cas9 system was used to perform genome editing for the knockout of p53 in WT-NSCs. Two different guide RNA (gRNA) sequences targeting the second exon (named gp53-1) and the fourth exon (named gp53-2) were designed. The site of gRNA is shown by the arrowheads, and the gRNA sequence is shown in the box below (Fig 1A). NSCs were obtained from embryonic day 14.5 (E14.5) mice brain [17]. Six days after culture, the p53 KO vectors were transfected. Three days after transfection, HRasL61 was transfected with GFP (NSCL61). GFP^+^ NSCL61 was sorted using a fluorescence-active cell sorter, and the cells were analyzed (Fig 1B). The p53 genotype in CTRL-NSCL61 and gp53-NSCL61 was analyzed by PCR (Fig 1C). In gp53-NSCL61, both the WT p53 genome and the edited p53 genome were detected (Fig 1C asterisk). When establishing SV40LT-NSCL61, overexpression of Simian virus 40 large T antigen was initially induced, and the cells were subsequently transfected with HRasL61 [19]. Because active HRas believed to induce cellular senescence, it was unexpected that gp53-NSCL61 contained the non-edited genome. Therefore, cell proliferation was analyzed. The numbers of CTRL-NSCL61 and gp53-NSCL61 were counted for 4 weeks and were found to be similar between gp53-NSCL61 and CTRL-NSCL61 in half medium (Fig 1D). These data suggest that several HRasL61-transfected NSCs undergo induced, senescence. However, the number of some cells was increased; therefore, these experiments required the use of CTRL-NSCL61.

**Fig 1.**
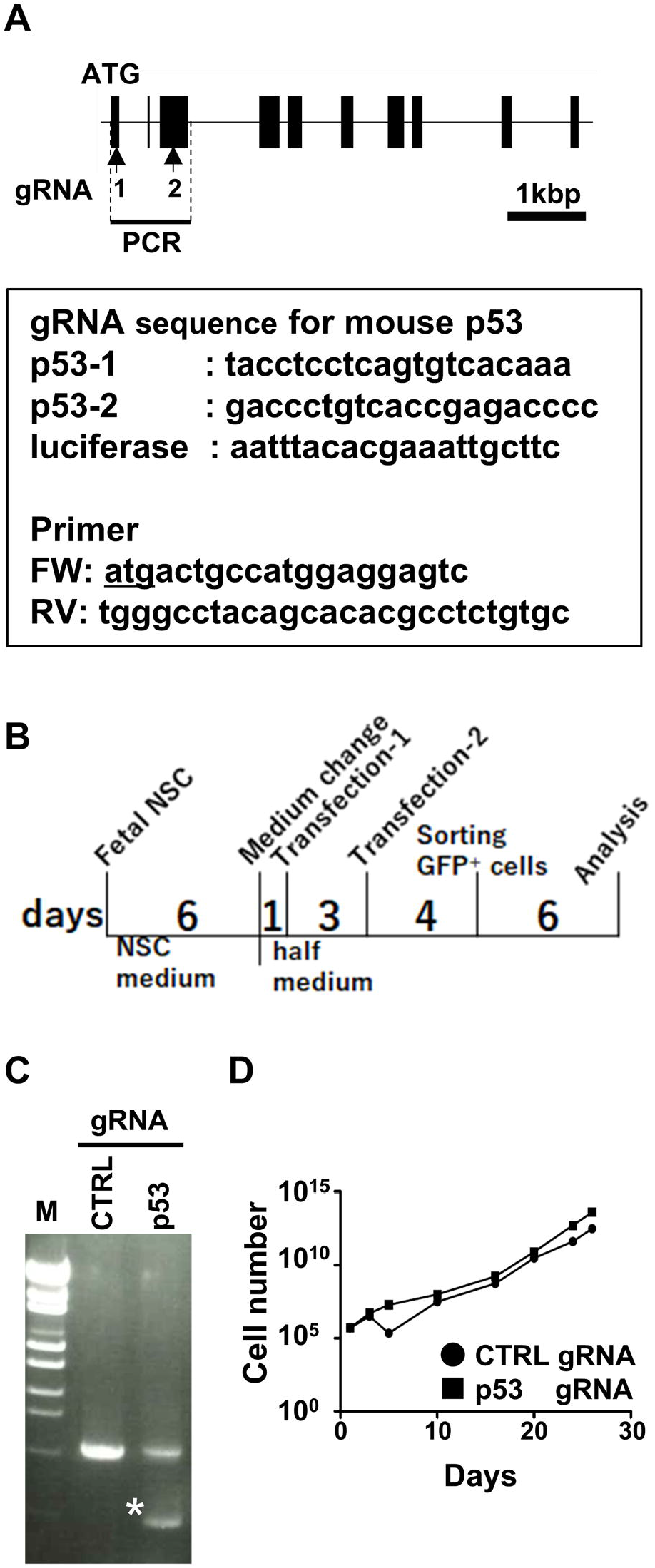
p53 knockout with the CRISPR/Cas9 system. (A) Structure of the mouse p53 gene. The mouse p53 gene contains 11 exons and 10 introns. The translation start codon (ATG) is located on the second exon. The guide RNA sequences for p53 and for luciferase (as a control) are shown in the box. (B) Schematic of the cell culture of vector transfection. (C) p53 genotype in CTRL- and in gp53-NSCL61. Luciferase gRNA was used as a negative control. The asterisk shows the edited genome. (D) Cell proliferation of gRNA- and HRasL61-transfected NSC. The cell numbers were counted for 4 weeks.

### Heterogeneous GICs generate tumors outside the brain

The tumorigenicity of the established gp53-NSCL61 was analyzed. A total of 100,000 CTRL-NSCL61s and gp53-NSCL61s were transplanted into the brain of NOD-SCID mice. Three weeks after transplantation, tumors were generated in gp53-NSCL61-transplanted mice (Fig 2A). The generated tumors were fused with skull and swollen (Fig 2A, right panel). The skulls were removed, and the brains were examined. Tumors were removed along with the skull, and a brain crushed by the tumor was detected (Fig 2B right panel, dashed circle). Penetration of gp53-NSCL61 was not found in the brain. The WT and edited genomes of p53 was detected in generated tumor (Fig 2C). Previous studies have shown that NSCL61s penetrated the brain [2,15,19,20], causing difference in the characteristics observed between gp53-NSCL61 with edited p53 gene and the normal p53 gene. gp53-NSCs in which the p53 gene had been edited were selected, and those cloned were analyzed.

**Fig 2.**
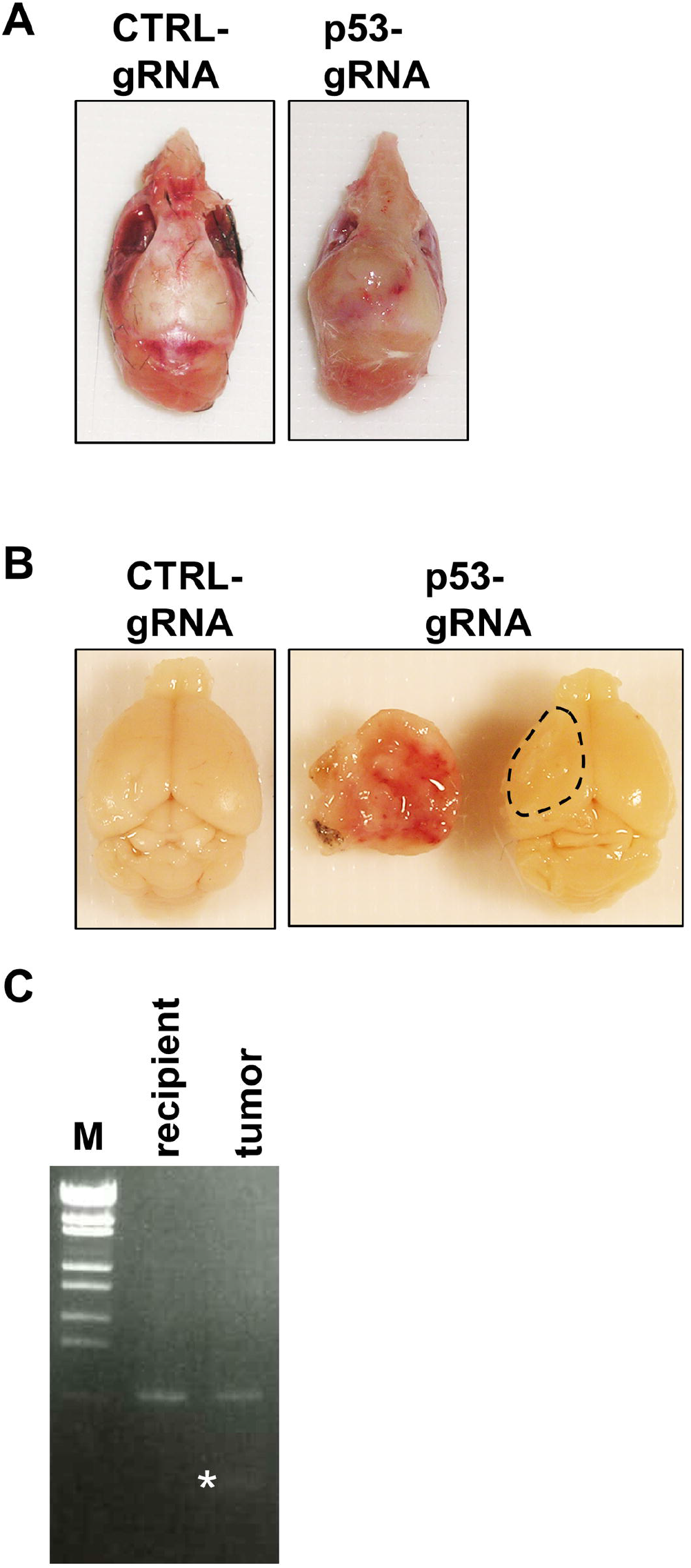
Tumorigenesis of gp53-NSCL61 bulk. CTLR-NSCL61 and gp53-NSCL61 were transplanted into the brains of NOD-SCID mice. Three weeks after transplantation, tumor was generated. (A) The skull of a NSCL61s-transplanted mouse containing the brain is showed. (B) NSCL61s-transplanted brains and tumors. The dashed line shows the contact surface of the brain and the tumor (n=9). (C) Genotype of p53 in generated tumor and recipient mice. The asterisk shows the edited p53.

### Cloning of p53 deficient NSCL61

Single-cell culture was performed to select only p53-edited gp53-NSCL61. The characteristics of the gp53-NSCL61 clone are showed in Fig 3. Genotyping PCR showed that only the p53-edited genome was detected in the gp53-NSCL61 clone (Fig 3A, asterisk). Western blotting analysis showed that the p53 protein detected in CTRL was not detected in gp53-NSCL61 (Fig 3B). These data suggest that p53 was not expressed in the gp53-NSCL61 clone.

**Fig 3.**
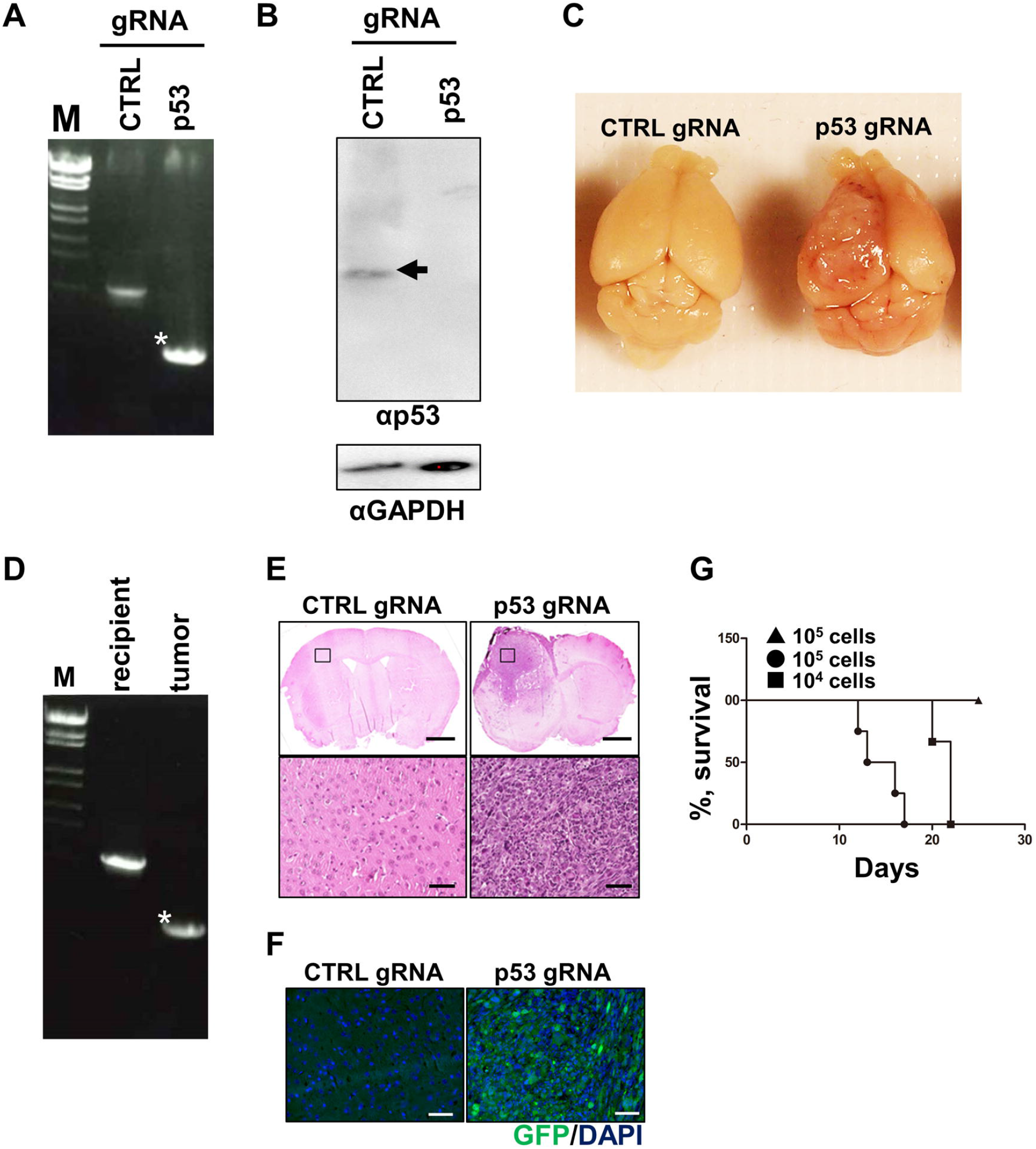
Tumorigenesis of the gp53-NSCL61 clone. (A) Genotype of CTRL-NSCL61 and gp53-NSCL61. The asterisk shows the edited genome. (B) The expression of p53 in CTRL-NSCL61 and gp53-NSCL61 was analyzed using Western blotting. GAPDH was detected as internal control. The allow shows the p53 signal. (C) NSCL61s-transplanted brains. NSCL61s were transplanted to NOD-SCID mice. Two weeks after transplantation, the brains were examined. (D) The p53 genotype in recipient mice and in generated tumors. The asterisk shows the edited genome. (E) Morphology of the generated tumors in the brains. The upper panels show paraffin sections stained with Hematoxylin-Eosin. The dashed circle shows tumor. The lower panels show high-magnification images. Scale bar: 1 mm (upper panels), 50μm (lower panels). (F) Sections were stained with anti-GFP antibody and DAPI. Scale bar: 50μm. (G) Survival curves for mice (n=4 for each line) injected with 10^4^ gp53-NSCL61 clone (■), 10^5^ gp53-NSCL61 clone (•), and CTRL-NSCL61 (▲).

### p53-deficient clones generate tumors in the brain

CTRL- and gp53-NSCL61 were transplanted into the brain of NOD-SCID mice. Two weeks after transplantation, the brains were observed. Tumorigenesis was detected only in the brain of gp53-NSCL61-transplanted mice, and the tumor penetrated the brain (Fig 3C). Genotyping PCR of the generated tumor showed that the p53 gene was edited (Fig 3D, asterisk). These data suggest that the generated tumors were derived from the transplanted gp53-NSCL61 clone. Paraffin sections of the NSCL61 transplanted brains were made and stained with hematoxylin-eosin for pathological analysis of tumors. High-magnification images showed hypercellularity, mitosis, nuclear pleomorphism, microvascular proliferation, and hemorrhage in the brain of gp53-NSCL61-transplanted mice. In contrast, these characteristics were not detected in CTRL-NSCL61-transplanted mice (Fig 3E, lower panels). These pathological features are similar to those of human GBM [4]. The origin of the formed tumors was also analyzed through immunostaining. Because established NSCL61s express GFP together with HRasL61, GFP was detected using immunohistochemistry. The black box in Fig 3E indicated that GFP was detected in the brain of the gp53-NSCL61 clone transplanted mice (Fig. 3F). This result also indicates that the formed tumors were derived from the transplanted gp53-NSCL61 clone. Survival time analysis demonstrated that 10^4^ and 10^5^ gp53-NSCL61 clone-transplanted mice expired approximately 3 and 2 weeks after transplantation (Fig 3G). 10^5^ CTRL-NSCL61 clone-transplanted mice survive for at least a 1 month. These dates suggest that gp53-NSCL61 clone is iGIC.

### The gp53-NSCL61 clone expresses GBM-specific genes

Our previous studies have shown that there are several GIC specific genes. Ten days after the forced expression of HRasL61, NSCL61s expressed an NSC-related marker gene (SOX2) and GIC marker genes (CD15, also termed stage-spesific embryonyc antigen-1 SSEA1; carcinoembryonic antigen-rerated cell adhesion molecule 1 [Ceacam1], also termed CD66a; and epitherial V-like antigen 1 [Eva1], also termed myelin protein zero like 2). Moderate expression levels of these GIC marker genes were observed in gp53-NSCL61 bulk (Fig 4). Regarding the gp53-NSCL61 clone, Eva1and CXeacam1 were expressed 10-fold and 20-fold higher, respectively (FIG 4). These four genes were expressed at level as similar to that measured in previously generated p53 KO-NSCL61s [14,15]. These results indicate that p53 was knocked out through genome editing, and WT-NSCs were converted to GICs.

**Fig 4.**
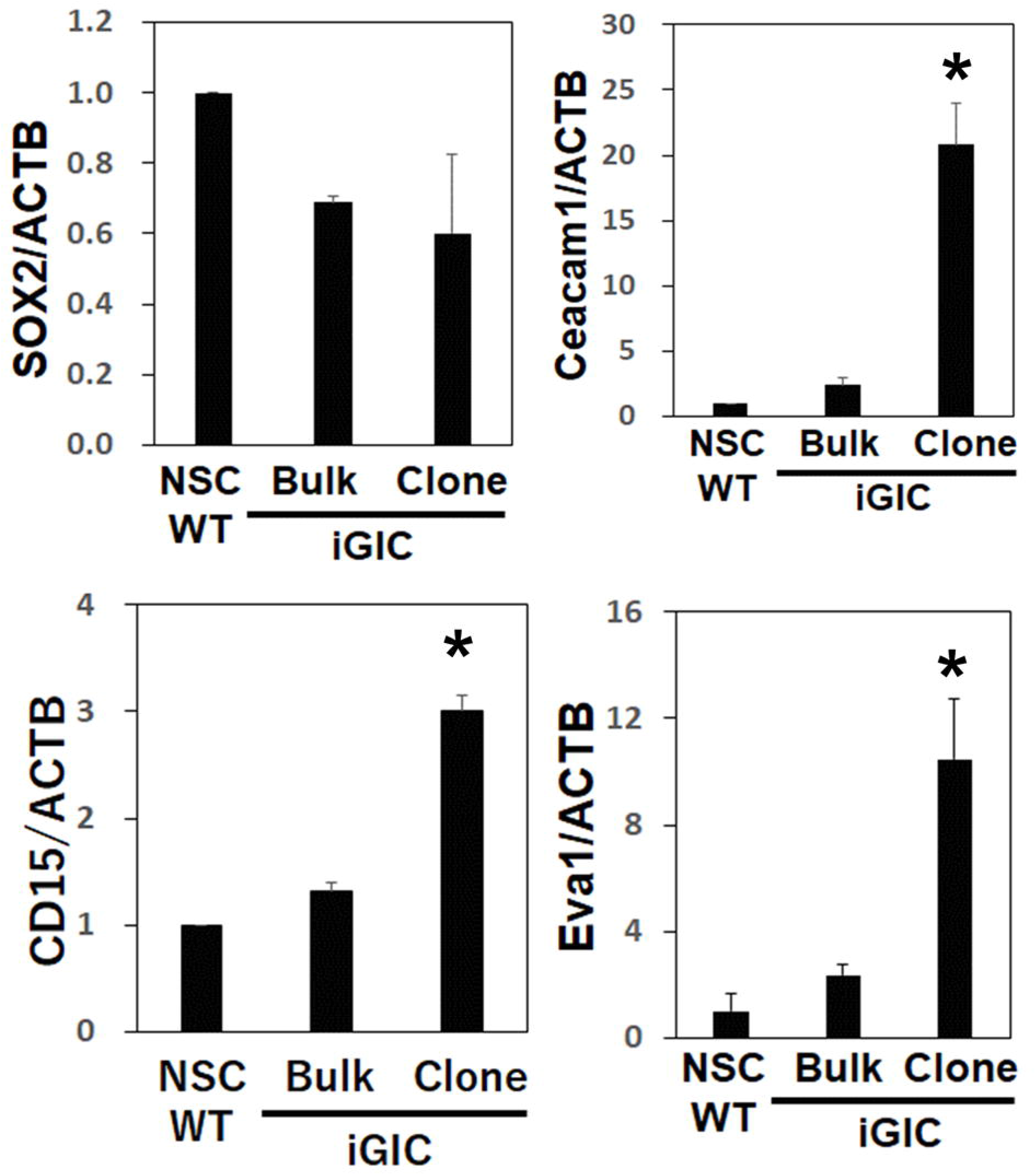
Expression of NSC and GIC-related genes. The expression levels of GIC-related genes (SOX2, CD15, Eva1, and Ceacam1) in WT-NSC, gp53-NSC61 bulk, and the gp53-NSCL61 clone were analyzed by qPCR. *p<0.05

## Discussion

In previous studies, iGIC were established by activating HRas in the NSCs of fetal p53 null mice, or adult Arf/INK4a null mice [2,13]. This study showed that TICs and GICs was induced from the WT mouse fetal NSCs by the knockout of the p53 gene through genome editing using the CRISPR/Cas9 system. The iGIC derived from fetal NSCs of esophageal cancer-related gene 4 (Ecrg4) null mice formed tumors in wild-type mice brain (Fig. 5) [20], However, the iGICs established this study did not form tumors when transplanted into WT mice. iGICs are induced from the adult brains of Arf/INK4a null mice [13]. However, tumorigenesis has not been confirmed for iGICs generated from adult p53 KO mouse brains. Genome editing enabled us to establish iGICs without using KO mice. The knockout of Ecrg4 together with p53 using genomic editing may generate iGICs that form tumors even in WT mice.

**Fig 5.**
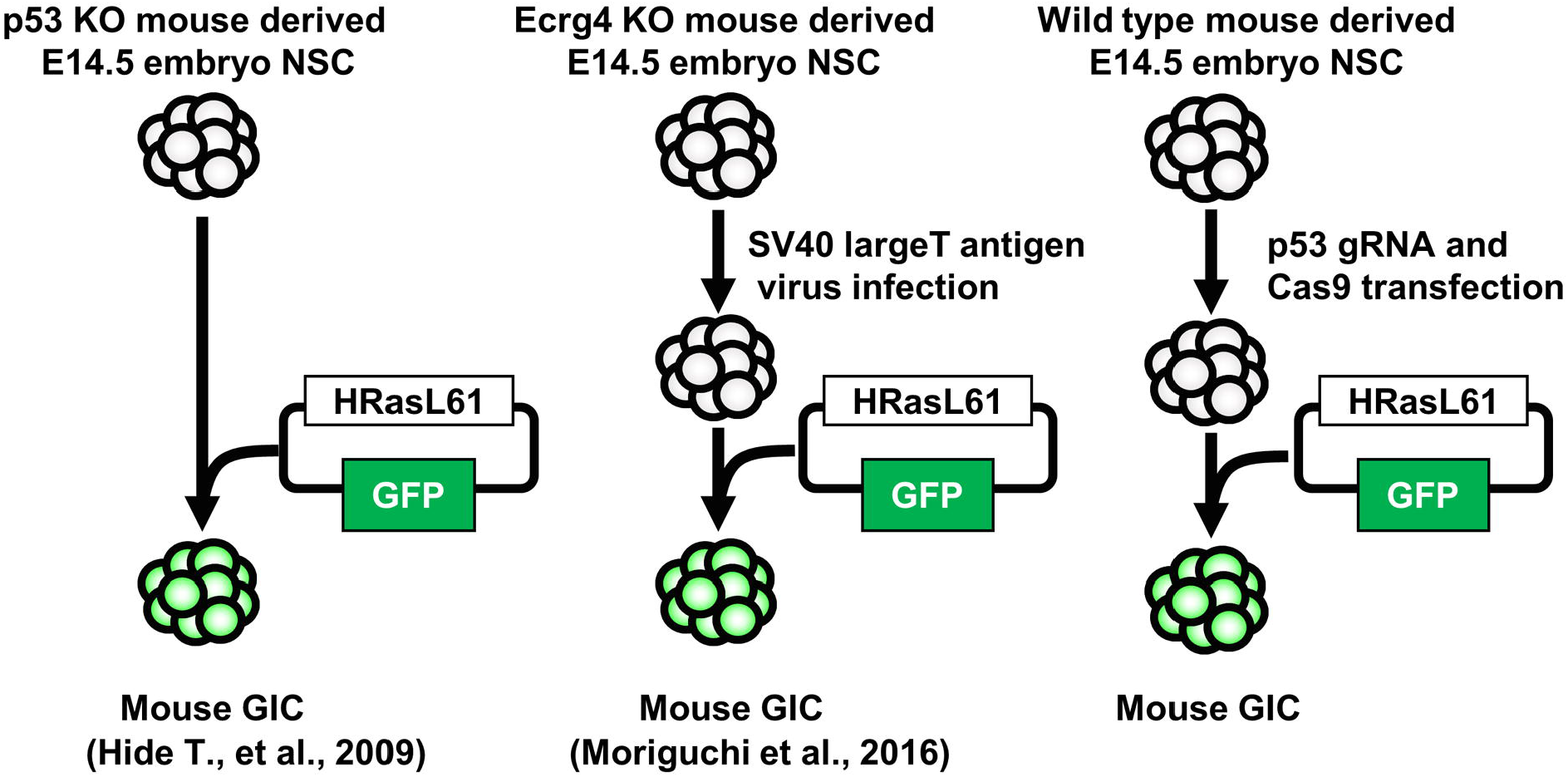
Schematic of the system used for the establishment of induced glioma-initiating cells.

It was possible to establish two patterns of TIC line (i.e., invasive and noninvasive). Because glioma (especially GBM) possesses the property of invasion into the brain, the gp53-NSCL61 bulk is iTIC, and it can be concluded that the gp53-NSCL61 clone is an iGIC. The TICs derived tumor formed contained p53 in the WT and deletion forms; therefor, cells with WT p53 may have somehow inhibited GBM invasion and malignant transformation. Interaction with the surrounding cells is necessary for the formation of GBM [21]. In our previous study, GBM cooperates with retinoic acid-related orphan receptor γ-positive T cells to suppress immune function [15]. It has been demonstrated that adjacent macrophages act as tumor-associated macrophages, thus GBM growth [14]. It is possible that p53-positive cells inhibited glioma tumorigenesis and invasion by directly or indirectly inhibiting such cell-cell interactions. In addition, when attempting to establish the GIC strain from a GBM tissue excised from a patient, a tumor could not be formed in the brain after repeated culture. Lowering the proportion of GICs among mutant cells may also inhibit tumor invasion. By analyzing the differences in the characteristics of TIC and GIC produced, it is possible to elucidate the mechanism of malignancy and to identify an approach in which cells contained in TIC inhibit the conversion of TICs to GICs.

In the TICs and GICs produced in this study, significant differences were detected in the expression levels of the existing GIC markers CD15 and the GIC-specific genes Ceacam1 and Eva1. However, no significant differences were detected for SOX2. Especially in Low grade glioma, high expression levels of Ceacam1 and Eva1 tend to shorten the survival time (S2 Fig). As the difference in the expression level of GIC markers between TIC and GIC is large, it has been suggested that they became standard for the establishment of GIC and the glioma grade.

The results of this study also confirm that p53 loss of function and HRas activation cause brain tumors to become malignant. Therefore, the development of new treatments targeting activation of p53 activation and suppression of HRas through genome editing is warranted.

## Supporting information

Saporting information

## Acknowledgements

The author thanks Dr. T. Kondo for providing pCMS-HRasL61 and the laboratory as well as Mr. Z. Wang and Ms. E. Ishizaki for technical assistance. This work was supported by JSPS KAKENHI Grant Number 18K07317.

## References

1. Singh SK, Clarke ID, Terasaki M, Bonn VE, Hawkins C, et al. Identification of a cancer stem cell in human brain tumors. Cancer Res. 2003; 63: 5821–5828.

2. Hide T, Takezaki T, Nakatani Y, Nakamura H, Kuratsu J, et al. Sox11 prevents tumorigenesis of glioma-initiating cells by inducing neuronal differentiation. Cancer Res. 2009; 69: 7953–7959.

3. Takanaga H, Tsuchida-Straeten N, Nishide K, Watanabe A, Aburatani H, et al. Gli2 is a novel regulator of sox2 expression in telencephalic neuroepithelial cells. Stem Cells. 2009; 27: 165–174.

4. Takezaki T, Hide T, Takanaga H, Nakamura H, Kuratsu J, et al. Essential role of the Hedgehog signaling pathway in human glioma-initiating cells. Cancer Sci. 2011; 102: 1306–1312.

5. Yamashita D, Kondo T, Ohue S, Takahashi H, Ishikawa M, et al. miR340 suppresses the stem-like cell function of glioma-initiating cells by targeting tissue plasminogen activator. Cancer Res. 2015; 75: 1123–1133.

6. Wong DJ, Segal E, Chang HY Stemness, cancer and cancer stem cells. Cell Cycle. 2008; 7: 3622–3624.

7. Parsons DW, Jones S, Zhang X, Lin JC, Leary RJ, et al. An integrated genomic analysis of human glioblastoma multiforme. Science. 2008; 321: 1807–1812.

8. Comprehensive genomic characterization defines human glioblastoma genes and core pathways. Nature. 2008; 455: 1061–1068.

9. Jeuken J, van den Broecke C, Gijsen S, Boots-Sprenger S, Wesseling P RAS/RAF pathway activation in gliomas: the result of copy number gains rather than activating mutations. Acta Neuropathol. 2007; 114: 121–133.

10. Rasheed BK, McLendon RE, Herndon JE, Friedman HS, Friedman AH, et al. Alterations of the TP53 gene in human gliomas. Cancer Res. 1994; 54: 1324–1330.

11. Reilly KM, Loisel DA, Bronson RT, McLaughlin ME, Jacks T Nf1;Trp53 mutant mice develop glioblastoma with evidence of strain-specific effects. Nat Genet. 2000; 26: 109–113.

12. Bogler O, Huang HJ, Kleihues P, Cavenee WK The p53 gene and its role in human brain tumors. Glia. 1995; 15: 308–327.

13. Sampetrean O, Saga I, Nakanishi M, Sugihara E, Fukaya R, et al. Invasion precedes tumor mass formation in a malignant brain tumor model of genetically modified neural stem cells. Neoplasia. 2011; 13: 784–791.

14. Kaneko S, Nakatani Y, Takezaki T, Hide T, Yamashita D, et al. Ceacam1L Modulates STAT3 Signaling to Control the Proliferation of Glioblastoma-Initiating Cells. Cancer Res. 2015; 75: 4224–4234.

15. Ohtsu N, Nakatani Y, Yamashita D, Ohue S, Ohnishi T, et al. Eva1 Maintains the Stem-like Character of Glioblastoma-Initiating Cells by Activating the Noncanonical NF-kappaB Signaling Pathway. Cancer Res. 2016; 76: 171–181.

16. Sakuma T, Yamamoto T Acceleration of cancer science with genome editing and related technologies. Cancer Sci. 2018; 109: 3679–3685.

17. Nakashima K, Yanagisawa M, Arakawa H, Kimura N, Hisatsune T, et al. Synergistic signaling in fetal brain by STAT3-Smad1 complex bridged by p300.Science. 1999; 284: 479–482.

18. Katayama S, Moriguchi T, Ohtsu N, Kondo T A Powerful CRISPR/Cas9-Based Method for Targeted Transcriptional Activation. Angew Chem Int Ed Engl. 2016; 55: 6452–6456.

19. Nishide K, Nakatani Y, Kiyonari H, Kondo T Glioblastoma formation from cell population depleted of Prominin1-expressing cells. PLoS One. 2009; 4: e6869.

20. Moriguchi T, Kaneumi S, Takeda S, Enomoto K, Mishra SK, et al. Ecrg4 contributes to the anti-glioma immunosurveillance through type-I interferon signaling. Oncoimmunology. 2016; 5: e1242547.

21. Pietras A, Katz AM, Ekstrom EJ, Wee B, Halliday JJ, et al. Osteopontin-CD44 signaling in the glioma perivascular niche enhances cancer stem cell phenotypes and promotes aggressive tumor growth. Cell Stem Cell. 2014; 14: 357–369

