## Supplementary figures and images for "Induction of tumor-initiating cells and glioma-initiating cells from fetal neural stem cells through p53 genome editing"

### Figure S1

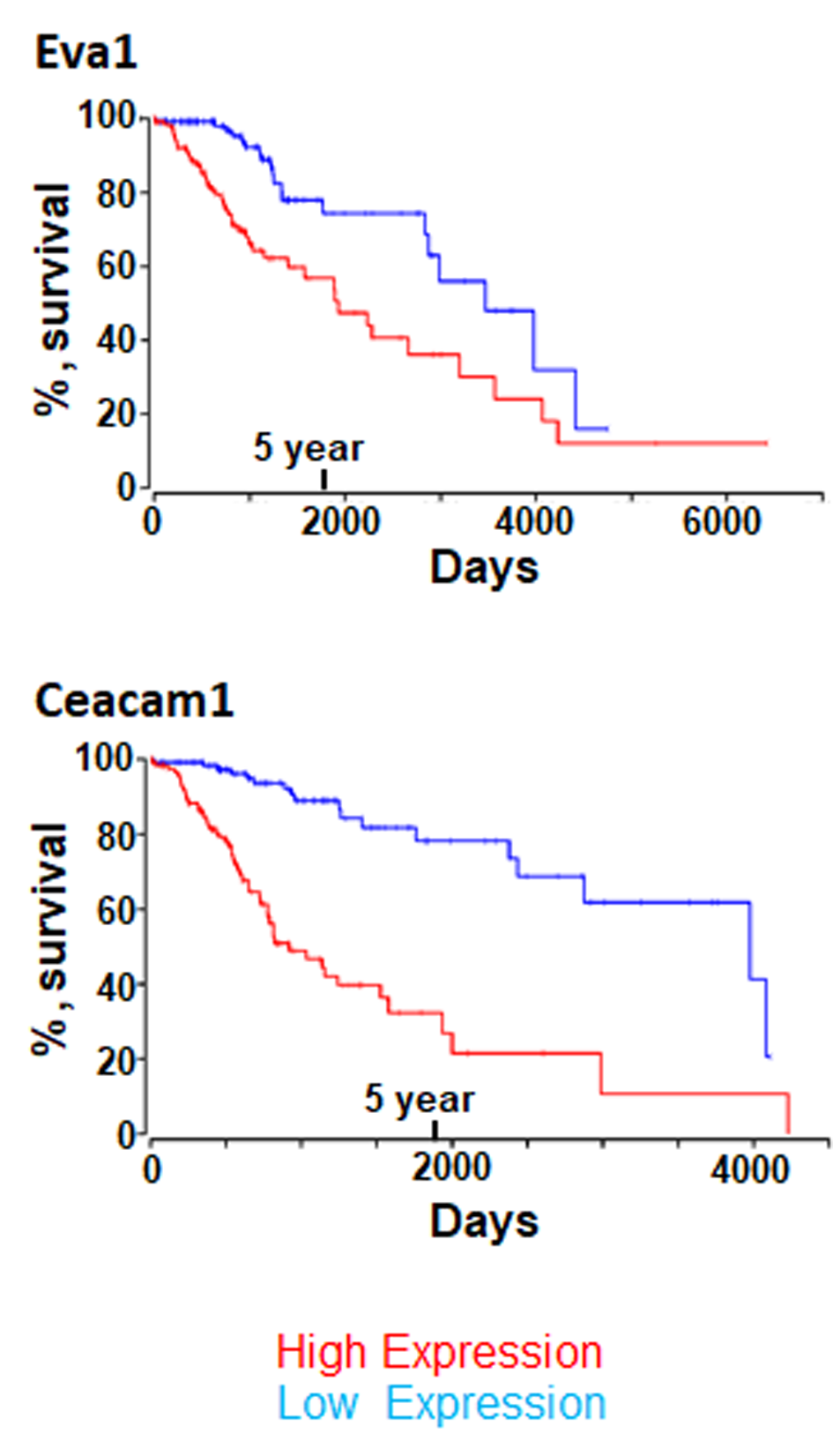

### Figure S2

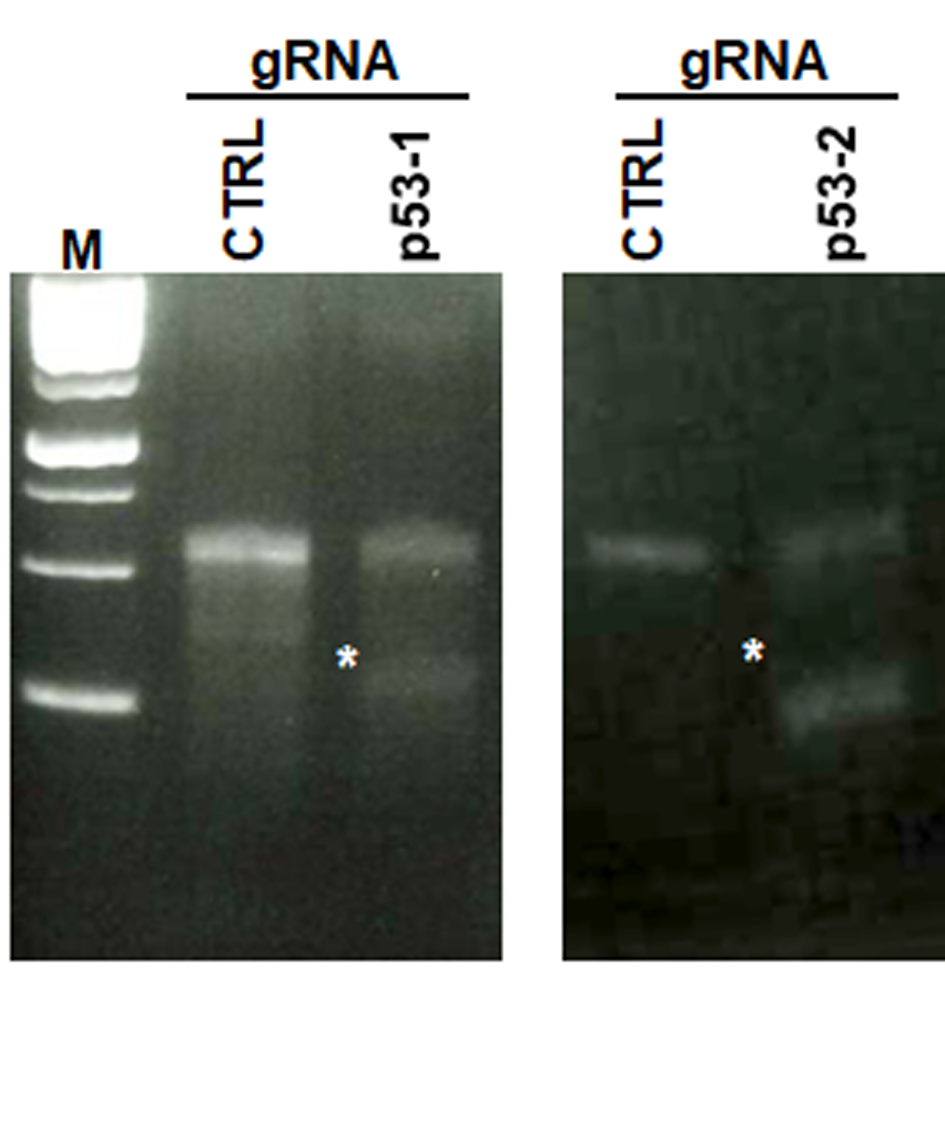
