## Supplementary material for "Induction of tumor-initiating cells and glioma-initiating cells from fetal neural stem cells through p53 genome editing": Saporting information

**Supporting information**

**S1 Fig. p53 knockout with the CRISPR/Cas9 system.** T7 Endonuclease 1 (T7E1) assay revealing genome editing at the p53-1 gRNA (A), and p53-2 sgRNA (B) in gp53-NSCL61 bulk. Asterisks indicated indel formation.

**S2 Fig. Survival time of Eva1 or Ceacam1 expressing glioma patient.** Clinical data from the OncoLnc database showed survival time of low-grade glioma patients with high expression levels of eva1 mRNA or ceacam1 mRNA (red line, Z-score> 2).

**S1 Table. Oligonucleotide sequences**

| Oligonucleotide | Sequences (5' to 3') | Applications |
| --- | --- | --- |
| p53 gRNA1-FW | cacctgccatggaggagtcacagt | gRNA |
| p53 gRNA1-RV | aaacactgtgactcctccatggca | gRNA |
| p53 gRNA2-FW | caccgaccctgtcaccgagacccc | gRNA |
| p53 gRNA2-RV | aaacggggtctcggtgacagggtc | gRNA |
| luciferase gRNA-FW | caccaatttacacgaaattgcttc | gRNA |
| luciferase gRNA-RV | aaacgaagcaatttcgtgtaaatt | gRNA |

**S2 Table. Primer sequences**

| Primer | Sequences (5' to 3') | Applications |
| --- | --- | --- |
| mp53-FW2 | atgactgccatggaggagtc | Gemomic PCR |
| mp53check-RV1 | tgggcctacagcacacgcctctgtgc | Gemomic PCR |
| mp53T7-FW1 | acttgcttataacttaatatcc | T7E1 assay |
| mp53T7-RV1 | ttgctatgtagtcaatgatgacc | T7E1 assay |
| mp53T7-FW2 | tggtaaggcccagagcagaaagg | T7E1 assay |
| mp53T7-RV2 | agtctacaggctgaagaggaacc | T7E1 assay |

**S3 Table. Primer sequences**

| Primer | Sequences (5' to 3') | Applications |
| --- | --- | --- |
| mSOX2-FW | aacggcagctacagcatgatgc | qPCR |
| mSOX2-RV | cgagctggtcatggagttgtac | qPCR |
| mCD15-FW | gattgcagcctgcgcttcaaca | qPCR |
| mCD15-RV | agtcgtggagttccttcaccag | qPCR |
| mEva1 qPCR-FW | tgacgtggaatttccgacctcg | qPCR |
| mEva1 qPCR-RV | agtgctggaagaggactacc | qPCR |
| mCeacam1 qPCR-FW | gaatccagtcagcgtcaggag | qPCR |
| mCeacam1 qPCR-RV | ccgccagacttcctggaatag | qPCR |
| ACTB qPCR-FW | agaagagctatgagctgcctgacg | qPCR |
| ACTB qPCR-RV | tacttgcgctcaggaggagcaatg | qPCR |
